# Cell-state-specific mechanisms of 5-azacytidine response in myelodysplastic neoplasms

**DOI:** 10.64898/2026.04.26.720516

**Authors:** Golam Sarower Bhuyan, Feng Yan, Mary N. T. Nguyen, Hoi Man Chung, Xiaoheng Zou, Veena Gullapalli, Lachlin Vaughan, Olivia Stonehouse, Henry R. Hampton, Sylvie Shen, Peter Truong, Ruchira Dissanayake, Elaheh S. Ghodousi, Swapna Joshi, Forrest C. Koch, Fabio Zanini, Fatemeh Vafaee, Yuanhua Huang, Julie A.I. Thoms, Omid R. Faridani, Christopher J. Jolly, John E. Pimanda

**Author notes:** Correspondence: Christopher J Jolly, John E. Pimanda. Equal first authors (Golam Sarower Bhuyan, Feng Yan).

## Abstract

5-azacytidine improves haematopoiesis and delays leukaemic progression in myelodysplastic neoplasms, but responses vary and are complicated by clonal mosaicism and heterogenous demethylation. Using a novel single-cell pipeline (“SCIMETAR-seq”), we found 5-azacytidine induced clonally distinct differentiation responses, with nuclear DNA demethylation occurring primarily in cycling progenitors. Despite the absence of significant nuclear DNA demethylation, quiescent stem cells underwent transcriptional remodelling *in viv*o, accompanied by 5-azacytidine-induced C•G-to-G•C mutations in mitochondria.

---

Myelodysplastic neoplasms or syndromes (MDS) are clonal malignancies driven by somatic mutations in haematopoietic stem cells (HSCs), leading to impaired haematopoiesis, cytopenias, and increased likelihood of AML progression^1–5^. Chronic Myelomonocytic Leukaemia (CMML), characterized by accumulation of immature CD34^+^ blasts and monocytes, is an MDS overlap neoplasm ^6, 7^. Higher-risk MDS/CMML patients are treated with hypomethylating agents (HMAs), including azacitidine (AZA), leading to improved blood counts and delayed AML progression in some patients ^8^. Metabolized AZA undergoes S-phase-dependent DNA incorporation where it covalently traps DNA methyltransferases (DNMT), leading to DNMT degradation and global DNA hypomethylation ^9, 10^. Single-cell approaches tracking mutations, HMA-incorporation, and gene expression are required to resolve how clonal mosaicism and heterogenous HMA-incorporation impact clinical response ^11, 12^.

Single-cell methods including Smart-RRBS ^13, 14^ measure DNA methylation alongside gene expression. However, requirement for cytosine deamination to measure DNA methylation limits tracking of C to T mutations, while needing to separate nuclei from cytoplasm/cDNA, limits throughput. Microfluidic platforms can mitigate the latter ^15^, while methylation-sensitive restriction enzymes enable limited analysis without eroding genotyping potential ^16^. Demethylation of endogenous viral elements is a robust proxy for genome-wide HMA-incorporation ^17^ that has been used in a single-cell bisulfite pipeline ^18^. Here we devised a bisulfite-free single-tube protocol-“SCIMETAR-seq” tracking single-cell Long-Interspersed Nucleotide Element 1 (LINE-1; L1) demethylation, immunophenotype, driver mutations, and full-length transcriptomes and used it to show that in haematopoietic stem and progenitor cells (HSPCs) cells undergoing AZA therapy L1 demethylation is associated with erythroid differentiation of common myeloid progenitors (CMP), but not with transcriptomic reprogramming of quiescent HSC and multipotent progenitors (MPP).

CpGs in L1 elements are hypermethylated in most somatic cells, and L1 demethylation can be used as a proxy for HMA-induced genome demethylation ^17, 19^. *Hpa*II-dependent End-Specific (ES) qPCR targeting a pair of human L1 5’UTR CpGs has been used to quantify L1 demethylation ^17^ since even one ^me^C in either strand of the target sequence (CCGG) blocks digestion, and only *Hpa*II-digested DNA generates a demethylation ES-qPCR signal (Extended Fig.1a,b). The human genome contains ∼146-290 full-length L1 ^20, 21^, so L1-ES-qPCR can detect demethylation in single-cell quantities of gDNA ^17^. We modified original L1-ES-qPCR oligonucleotides to improve homology to a full-length L1 consensus^20, 21^ and to convert the internal reference ES-qPCR oligonucleotides into conventional qPCR primers (Extended Fig.1a,b). These increased L1-ES-qPCR sensitivity, whilst retaining specificity for *Hpa*II digestion to generate demethylation (FAM), but not internal reference (HEX) qPCR signals (Extended Fig.1c,d). Demethylation and reference qPCRs exhibited similar efficiencies (*E*=1.861.88; Extended Fig.1e), allowing the fold-increase in demethylation in DNA sample *x,* relative to control (i.e. HMA-free) conditions, to be approximated as 1.87^ΔΔCt^ (Extended Fig.1f).

We validated single-cell L1-ES-qPCR using the human MDS cell line, MDS-L (Extended Fig.1g). LC-MS ^22^ detected similar cytidine demethylation levels in response to 0.3 or 1.0 µM AZA (Extended Fig.1h). L1-ES-qPCR detected dose-dependent demethylation, although L1-demethylation was less than the theoretical limit estimated by replacing *Hpa*II with *Msp*I (Extended Fig.1i, left); *Msp*I cleaves CCGG sites regardless of DNA methylation ^23^ (Extended Fig.1a). L1-ES-qPCR of single MDS-L cells produced results comparable to bulk MDS-L L1-ES-qPCR data (Extended Fig.1i, right). L1-ES-qPCR was integrated into SCIMETAR-seq (Fig.1a).

**Figure 1:**
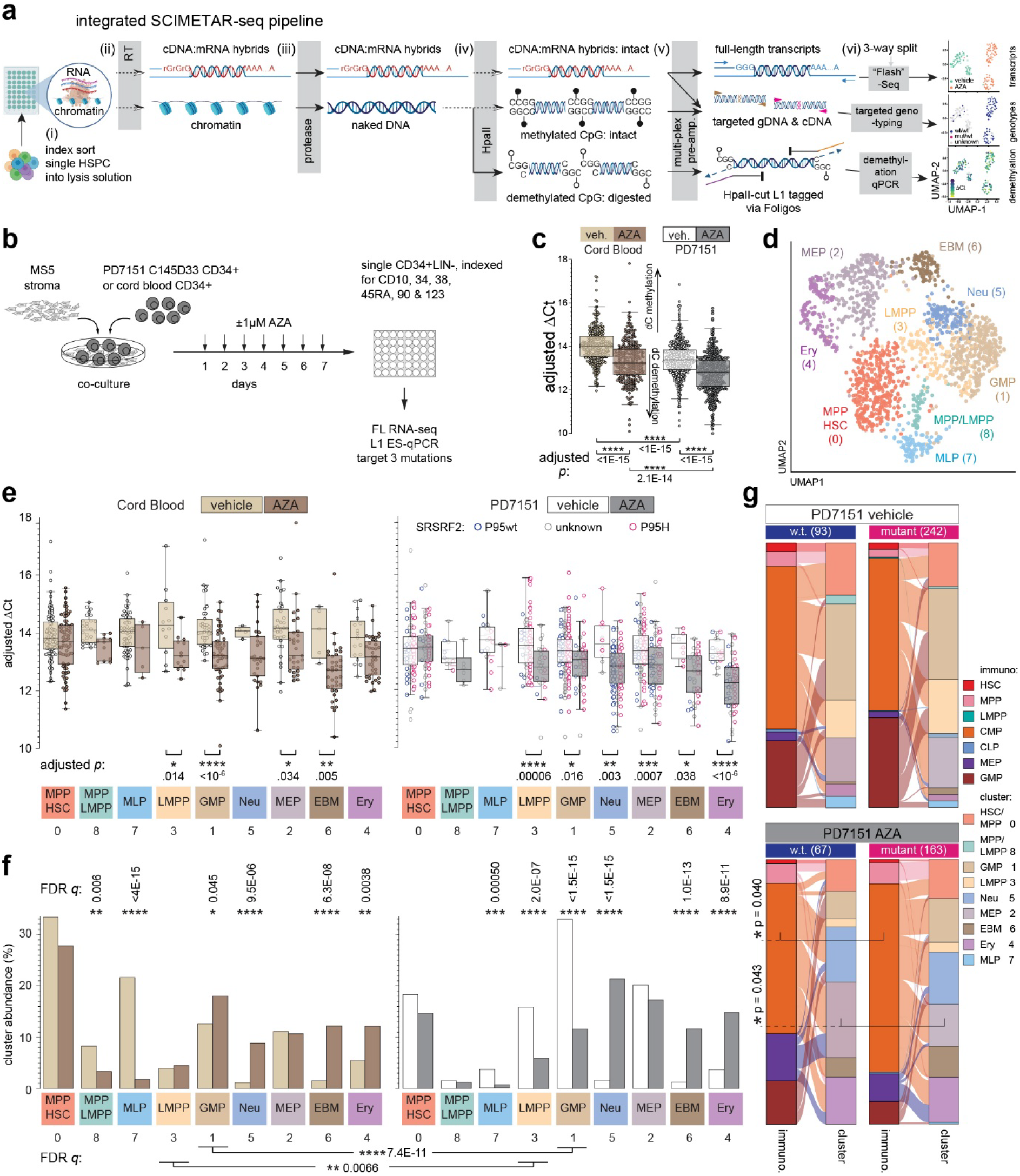
SCIMETAR-seq analysis of *in vitro* AZA-treated cells. **(a)** SCIMETAR-seq pipeline. Components of this panel were created with BioRender.com. **(b)** *in vitro* AZA co-culture experiment. **(c)** Plate-normalised L1-ES-qPCR ΔCt for cells passing QC; *p:* Šídák’s multiple comparisons test. **(d)** UMAP embedding of CB and PD7151 cells, coloured by cluster. **(e)** Per-cluster single-cell adjusted ΔCt; *p:* Šídák’s multiple vehicle vs AZA comparisons. PD7151 *SRSF2*-genotypes: wt (blue), p.P95H (pink). **(f)** Relative abundance per cluster; *q:* multiple 2×2 contingency tests. **(g)** Sankey plots linking immunophenotypic and transcriptomic identities of PD7151 cells; Unadjusted *p:* single 2×2 contingency tests.

SCIMETAR-seq was piloted on 186 single MDS-L cells ±1 µM AZA (Extended Fig.2a) with recovery of up to 7.5×10^3^ genes per cell with full-length mRNA coverage (Extended Fig.2b,c). High coverage cells (>1000 genes recovered; 87% of wells) formed four transcriptomic clusters (Extended Fig.2d) with varying mean L1-ES-qPCR ΔCt scores (Extended Fig.2e,f). Differentially expressed genes were enriched for mitochondrial and protein translation GO pathways (Extended Fig.2g). Targeted *TET2* exon 3, *TET2* exon 6, and *SRSF2* regions (target genotyped previously ^24^), were adequately recovered (≥10 reads) from 100%, 100% and 51% of single MDS-L cells respectively (Extended Fig.2h).

We applied SCIMETAR-seq to HSPCs from cord blood (CB) or a CMML patient, PD7151 ^24, 25^ (Extended Fig.3a,b). HSPCs were AZA-treated *in vitro* then index sorted for SCIMETAR-seq (Fig.1b, Extended Fig.3c). AZA treatment increased immunophenotypic CMP and megakaryocyte erythroid progenitors (MEP) at the expense of granulocyte monocyte progenitors (GMP) (Extended Fig.3d). L1 methylation in AZA-treated cells was lower than vehicle controls as expected, and lower in PD7151 compared to CB (Fig.1c), due to prior exposure of PD7151 cells to AZA *in vivo* (Extended Fig.3a). Shared enriched pathways included chromosome segregation and myeloid homeostasis in the most AZA-demethylated cells, and cytoplasmic translation, cell-cell adhesion and MHC class II antigen presentation in the least AZA-demethylated cells (Extended Fig.3e). Significant L1 demethylation in response to AZA was restricted to progenitors (CMP, MEP and GMP; Extended Fig.3f).

Nine transcriptomic clusters (Fig.1d, Extended Fig.3g), were annotated with reference to ^26^ (Extended Fig.3h). Except for HSC, MPP and multi lymphoid progenitor (MLP), cell surface immunophenotype poorly predicted RNAseq-based phenotypes (Extended Fig.3i). AZA did not significantly demethylate L1 in cluster-0 HSC/MPP (Fig.1e) or alter HSC/MPP frequencies (Fig.1f), likely due to low AZA incorporation in quiescent cells. AZA increased frequencies of granulocyte- and erythrocyte-committed RNAseq clusters (5,6,4) at the expense of lymphoid-biased clusters (8,7; Fig.1f), accompanied by substantial demethylation within the expanded myeloid clusters (Fig.1e).

Genotyping for *TET2* exon 3, *TET2* exon 6 and *SRSF2* mutations in PD7151 passed QC (≥10 relevant reads) in 99%, 96% and 47% of cells, respectively (Extended Fig.3c, j). 1392/1431 PD7151 cells genotyped carried exon 3 and 6 *TET2* mutations (in separate alleles), and 72% of these carried the *SRSF2* mutation; no mutations were detected in CB (Extended Fig.3j). Overall L1 demethylation did not alter with *SRSF2* mutation status (Fig.1e). However, AZA induced differentiation from CMP into cluster-2 MEP in *SRSF2*-wt cells more than in *SRSF2*-mutant cells (Extended Fig.3k, Fig.1g).

Finally, we applied SCIMETAR-seq to HSPC from PD7151 collected immediately before (day (D) 1) and after (D8) AZA treatment cycle (C) 4 (Fig.2a). Expanded MEP and reduced CMP frequencies were observed post-AZA-treatment (Extended Fig.4a). QC metrics were similar to previous, with reduced *SRSF2* genotyping in C4D1 (Extended Fig.4b). Mean L1 methylation levels were unchanged between D1 and D8 (Fig.2b). Nonetheless, transcriptomic differences were observed in the least versus most demethylated cells post-AZA (C4D8; Fig.2c); enriched pathways were comparable to *in vitro* AZA treatment (Extended Fig.3e). Enrichment for DNA recombination, nucleotide metabolism and mitochondrial gene expression in the most demethylated C4D8 cells aligns with known adaptations to acute AZA toxicity ^27–29^. Normalised ΔCt L1-ES-qPCR correlated with RNAseq C4D8 S-phase or G2M-phase scores (Fig.2d, Extended Fig.4c), and median L1-methylation was highest in HSCs (Extended Fig.4d).

**Figure 2:**
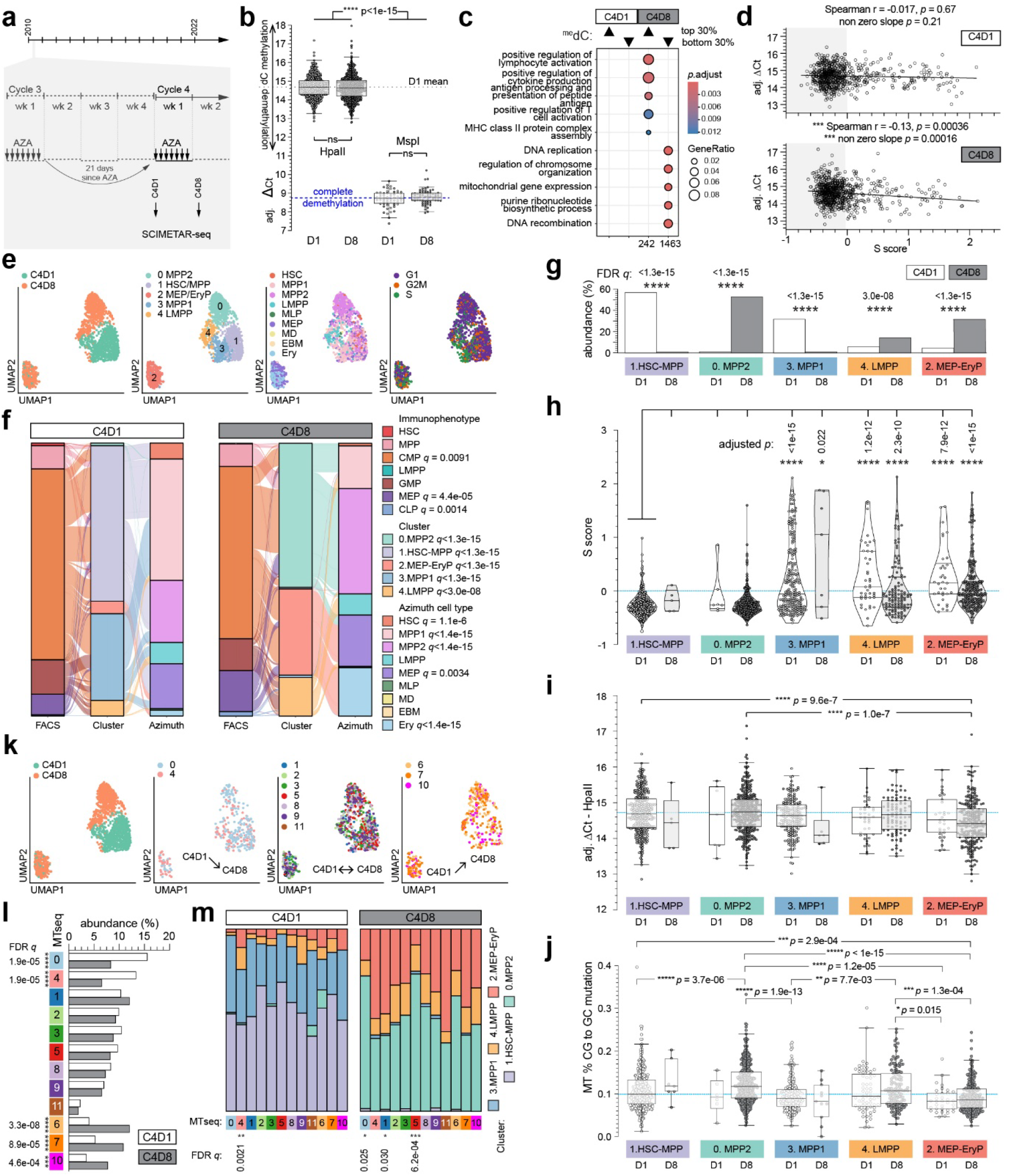
SCIMETAR-seq analysis of *in vivo* AZA-treated cells. **(a)** Clinical (*in vivo*) AZA treatment. **(b)** Adjusted L1-ES-qPCR ΔCt; Welch’s t-test. “Complete demethylation” = median *Msp*I-dependent ΔCt. **(c)** Pathway enrichment for genes differentially expressed between most (▴) or least (▾) methylated cells at C4D8. **(d)** ΔCt vs. transcriptomic S-score (shading = not S-phase); Spearman correlation and linear regression. **(e)** UMAP embedding of PD7151 cells colored by timepoint, cluster, reference transcriptome, or cell cycle. **(f)** Sankey plots linking cell identities; *q:* multiple contingency tests for D1 vs D8. **(g)** Cluster frequencies; *q:* multiple contingency tests for D1 vs D8. **(h)** S-score distributions; *p:* multiple nonparametric comparisons to C4D1 cluster-1. **(i)** Per-cluster single-cell adjusted ΔCt; *p:* Dunnett’s multiple comparisons. **(j)** Frequencies of C•G to G•C mutation in mt by day and cluster (blue line = C4D1 cluster-1 median); *p:* multiple nonparametric all-ways comparisons. **(k)** UMAP embedding of PD7151 C4 cells colored by shrinking, unaffected, or expanding mt lineages. **(l)** Distributions of mt lineages; *q:* multiple 2×2 contingency tests. **(m)** RNAseq cluster distributions for each mt lineage; *q:* multicomparison corrected 2×5 contingency tests for D1 (left) or D8 (right).

Pre- vs post-AZA cells were mostly transcriptionally distinct (Fig.2e-g), but close overlaps in cluster-2 suggest this was not merely a batch effect. Clusters-1 and -0 contained mostly quiescent MPP and HSC, while clusters-3 and -4 contained non-quiescent MPP, Lymphoid-Primed Multipotent Progenitor (LMPP), MEP and MLP, and cluster-2 contained non-quiescent MEP and erythroid progenitors (EryP) (Fig.2e-h). Haematopoiesis was highly clonal at this near-diagnosis timepoint, because 1219/1277 cells genotyped carried both exon 3 and 6 *TET2* mutations, but only 1/726 genotyped cells was classified as *SRSF2* mutant (Extended Fig.4e,3a).

L1 in cluster-2 and -3 significantly demethylated during AZA cycle 4 (Fig.2i). Cluster-2 (MEP/EryP) expanded whilst cluster-3 (erythroid-biased MPP1 ^26^) shrank (Fig.2f), suggesting that AZA-induced differentiation from cluster-3 to cluster-2 is likely the mechanistic basis of increased circulating haemoglobin and platelets following AZA cycle 6 (Supplementary Table 1). In contrast, equivalent L1 methylation in cluster-1 (mostly D1 HSC/MPP) compared to cluster-0 (mostly D8 HSC/MPP; Fig.2i) was consistent with minimal AZA incorporation in quiescent cells. Nonetheless, C4D8 cluster-0 cells were significantly enriched for NFkB activation compared to C4D1 cluster-1 cells (Extended Fig.4f), suggesting that quiescent HSC/MPP responded to inflammation during AZA cycle 4. We hypothesized that AZA incorporation into mitochondrial (mt) DNA might be an intrinsic source of inflammatory signaling in quiescent HSC. We tested this by quantifying C to G (also G to C) mutations in mt reads in scRNAseq data (AZA-incorporation into nuclear DNA is known to induce a C to G mutation signature ^30–32^). We uncovered a 3.8-fold increase (*p*<1e-15) in C to G mutation from 2010 (C4) to 2022 (C145) in PD7151 HSPC mt (Extended Fig.5a). Increased mutation frequency was also evident in the cord blood cells that had been cultured for seven days with AZA compared to vehicle (Extended Fig.5b); this confirmed AZA as the C to G mt mutation driver. Within treatment cycle 4, the highest increase in C to G mt mutation from D1 to D8 occurred in D8 cluster-0 cells compared to D1 cluster-1 cells (1.2-fold increase, *p=*3.7e-06; Fig.2j), despite cluster-0 cells having the lowest cell cycle scores of C4D8 clusters (Fig.2h). Thus, mt DNA replication in quiescent HSC/MPP over as little as one treatment cycle was sufficient for detectable AZA-induced mt mutation. We deduce that AZA-induced damage to mtDNA (*via* ring-opening of incorporated dAZA ^31^) would have increased the release of mtDNA to the cytosol to enhance inflammatory signaling in cluster-0 cells (Extended Fig.4f) ^33^.

We also observed elevated TGFβ-signaling and WNT-signaling in D8 cluster-0 compared to D1 cluster-1 cells (Extended Fig.4f), suggesting that quiescent HSC/MPP may have responded to external AZA-induced differentiation cues from the bone marrow microenvironment during treatment cycle 4.

Given the lack of *TET2* and *SRSF2* mosaicism in C4, we explored use of mt variants as lineage barcodes to track clones. SNPmanifold identified 12 mt lineages from the most abundant mtSNP (Extended Fig.5c). Frequencies of mt lineage-0 and -4 cells reduced significantly from C4D1 to C4D8, while mt lineages-6, -7 and -10 increased (Fig.2k, l). Furthermore, at C4D8 mt lineage-0 and -5 HSPCs were enriched in quiescent RNAseq-0 (MPP), while mt lineage-1 cells were enriched in non-quiescent erythroid-biased RNAseq-2; such biases were less apparent at C4D1 (Fig.2m).

In summary, we applied SCIMETAR-seq to track immunophenotype, demethylation, nuclear and mt mutations, and transcriptomes in primary patient samples, finding that AZA-incorporation into nuclear DNA of HSPCs cells is restricted to proliferating progenitors and unaffected by *SRSF2* genotype. Nonetheless, *SRSF2-* or mt-genotyping identified clones with distinct differentiation biases. Quiescent HSC/MPP, which incorporated little or no AZA into genomic DNA, responded to clinical AZA treatment, which was associated with increased intrinsic inflammation and AZA-incorporation into mtDNA.

## Acknowledgements

The authors thank staff and donors of the Sydney Cord Blood Bank for providing cord bloods for research. Some of the data presented in this work was acquired by personnel and/or instruments at the Mark Wainwright Analytical Centre (MWAC) of UNSW Sydney, which is in part funded by the Research Infrastructure Programme of UNSW. Funding for this work came from National Health and Medical Research Council (2011627 to HRH, ORF, FZ, CJJ, JEP), Anthony Rothe Memorial Trust (RG213236 to JAIT), Cancer Institute NSW (2022/TPG2152 to JAIT, JEP).

## Authorship contributions

GSB, VG, LV, HRH, SS, FZ, ORF, CJJ, JEP designed research; GSB, MNTN, XZ, VG, LV, OS, HRH, SS, PT, RD, ESG, SJ performed research; GSB, FY, OS, FCK, HMC, FV, YH, JAIT, CJJ,

JEP analyzed and interpreted data; GSB, FY, JAIT, CJJ, JEP wrote the manuscript.

## Conflict of interests

The authors declare no relevant conflicts of interest.

## Methods

### Sample sources and data acquisition

#### Cell lines

The human myelodysplastic neoplasm cell line MDS-L ^34^ was kindly provided by Dr. Kaoru Tohyama (Department of Laboratory Medicine, Kawasaki Medical School). Cells were maintained at 37 °C in a humidified incubator with 5% CO₂ and cultured in RPMI-1640 medium (Thermo Fisher Scientific, 11875-093) supplemented with 10% fetal bovine serum (Sigma-Aldrich, F9423), penicillin–streptomycin (Gibco, 15140-122; 100 units/mL), GlutaMAX (Gibco, 35050061; 2 mM), recombinant human interleukin-3 (25 ng/mL; Miltenyi Biotec, 130-095-069), and β-mercaptoethanol (50 µM; Sigma-Aldrich, M6250). Cell line identity was confirmed by short tandem repeat profiling performed at the Garvan Institute of Medical Research, and routine mycoplasma testing was performed at UNSW. MS-5 stromal cells ^35^ were provided by Dr Karen MacKenzie (Children’s Medical Research Institute, Westmead, NSW, Australia) and maintained at 37 °C in a humidified incubator with 5% CO₂ and cultured in αMEM medium (Thermo Fisher Scientific, 12571-063) supplemented with 10% fetal bovine serum (Sigma-Aldrich, F9423), penicillin–streptomycin (Gibco, 15140-122; 100 units/mL), and GlutaMAX (Gibco, 35050061; 2 mM).

#### Patient Samples

Peripheral blood and BM samples from a patient with CMML (patient PD7151/H198303, patient characteristics and variant allele frequencies (VAF) are described in earlier genomic studies ^24, 25^) were collected with written informed consent in accordance with the Declaration of Helsinki, under institutional ethics approvals 08/190 from the South Eastern Sydney Local Health District, NSW, Australia. Cord blood units were provided by Sydney Cord Blood Bank. Use of samples received ethical approval (2022_ETH00727) from the South Eastern Sydney Local Health District Human Research Ethics Committee. CD34^+^ HSPCs were isolated from BM and CB as previously described ^12^.

#### Co-culture and Azacitidine treatment

CD34⁺ HSPCs and MDS-L cells were cultured *in vitro* using the MS-5 stromal cell co-culture system ^25, 36, 37^. MS-5 cells were seeded at 1.5 × 10⁵ cells/mL in αMEM (Thermo Fisher Scientific, 12571-063) supplemented with 10% fetal bovine serum (Sigma-Aldrich, F9423-500 mL) and transferred to a humidified incubator at 37 °C supplemented with 5% CO₂ for 24 hours to establish a confluent monolayer. MDS-L cells were resuspended in MDS-L medium (see above), and overlaid onto MS-5 stromal monolayers. Azacitidine (Selleckchem, S1782; 0.3µM or 1.0µM final, or vehicle) was then added daily for 4 days, starting immediately, from single-use 0.1M stock stored in aliquots at -80C in anhydrous dimethyl sulfoxide vehicle (DMSO, Sigma-Aldrich, D2650). Primary CD34⁺ HSPCs were resuspended in Gartner’s medium containing IMDM (Thermo Fisher Scientific, 12440-053) with 12.5% fetal bovine serum (Sigma-Aldrich, F9423-500 mL), 12.5% horse serum (Thermo Fisher, 16050130), 2 mM L-glutamine (Thermo Fisher Scientific, 25030081), 1 µM hydrocortisone (Sigma-Aldrich, H0888-1G), 8 µg/mL monothioglycerol (Sigma-Aldrich, M6145-25 mL) and 50 µg/mL gentamycin (Thermo Fisher Scientific, 15750-060), then overlaid onto MS-5 stromal monolayers. Starting 48h later, AZA (1µM final, or vehicle) was added daily for 7 consecutive days.

#### Cell staining and sorting

Cells were harvested from MS-5 co-cultures using Trypsin-EDTA (Gibco, 15400054), pelleted by centrifugation (500g for 7 min at 20C), washed with Dulbecco’s phosphate-buffered saline (PBS, Thermo), and resuspended in FACS buffer (2% fetal bovine serum (Sigma-Aldrich F9423), 1 mM EDTA (Sigma-Aldrich 03690) in PBS). Cryopreserved PD7151 CD34^+^ HSPCs from clinical time points C4D1 and C4D8 (pre- and post-AZA during treatment cycle 4) were thawed, washed with PBS, and resuspended in FACS buffer. Samples were stained on ice for 30 min using a BM antibody panel (Supplementary Table 2), then washed once with FACS buffer. Sorting was performed at the UNSW Mark Wainwright Analytical Centre using a FACSAria III cell sorter (Becton Dickinson). Unstained, fluorescence-minus-one, and single-colour compensation controls were used to define gating and correct for spectral overlap. Single cells sorted for L1-ES-qPCR only were sorted into 96-well plates containing 4 µL/well lysis buffer (experiment specific). Three empty wells were included in each plate. Immediately after sorting, plates were sealed with foil (Biorad MSF1001B), centrifuged at 300g for 30 s at 4 °C, frozen on a bed of dry ice, then stored in liquid nitrogen until further processing.

#### Isolation of DNA and Cytidine LC-MS

Genomic DNA from bulk MDS-L cells was extracted using the All-in-One DNA/RNA Miniprep Kit (BioBasic, BS88203) for mass spectrometry (MS) experiments, or the AllPrep DNA/RNA Mini Kit (Qiagen, 80204) for L1-ES-qPCR analyses. DNA concentration was estimated using a NanoDrop spectrophotometer (Thermo Fisher Scientific) for MS or a Qubit fluorometer (Thermo Fisher Scientific) with the Quant-iT Qubit dsDNA HS Assay Kit (Invitrogen, Q32851) for L1-ES-qPCR. For single or small numbers of cells sorted into 96-well plates, DNA was liberated from pelleted cells by Qiagen protease (0.6 mg/mL) and heating to 50°C for 90 min, then 75°C for 30 min. 2′-deoxycytidine and 5-methyl-2′-deoxycytidine levels in genomic DNA were quantified using LC-MS as previously described ^22, 27^.

#### Primer design for target amplicons

Bulk variant allele frequency analysis identified two pathogenic mutations in *TET2* (hg19 chr4:106158367_AAGAC>A, p.K1090fs15; hg19 chr4:106164802_GC>G, p.A1224fs2) and one in *SRSF2* (hg19 chr17:74732959_G>T, p.P95H) in PD7151 ^24^ (see Extended Fig.3a). PCR primers flanking these mutation sites (Supplementary Table 3, Extended Fig.1k) were from previous work ^24^. Illumina-compatible universal amplicon barcoding oligos (Supplementary Table 4) incorporated degenerate bases and variable-length stagger sequences to aid cluster discrimination.

#### Rationale for using LINE-1 (L1) methylation as a proxy measure of AZA incorporation

Since endogenous viral elements (EVE) demethylate and increase transcription in response to HMA, potentially inducing *via* viral mimicry ^38–41^, and are present in very high copy numbers, demethylation of EVE is a useful lower cost proxy for HMA-incorporation into DNA ^17, 18^. CpGs located in L1 elements are hypermethylated in most somatic cells. Demethylation of L1, which comprise ∼17% of human DNA, can therefore be a useful proxy for whole genome demethylation in response to HMAs ^17, 19^. *Hpa*II-dependent End-Specific (ES) qPCR targeting a close-spaced pair of human L1 5’UTR CpGs is a novel approach to quantify L1 demethylation ^17^. The presence of even one ^me^C in either strand of the *Hpa*II target sequence CCGG blocks *Hpa*II digestion, and only *Hpa*II-digested DNA generates a demethylation ES-qPCR signal (Extended Fig.1a,b). Since there are about 146 to 290 full length L1 per human genome ^20, 21^, L1-ES-qPCR can detect demethylation in single-cell equivalent quantities of genomic DNA ^17^.

#### Bulk L1-ES-qPCR

“Original” reagent L1-ES-qPCR reactions were performed as described ^17^. “Updated” reagent bulk DNA ES-qPCR reactions (20 µL reactions in 96-well plates, including DNA from <300 cells per well) consisted of 50 mM KCl (Merck, 60142), 7 mM MgCl₂ (Merck, M1028), 200µM dNTP mix (Thermo Fisher Scientific, R0181), 0.5 U Hot Start Taq DNA polymerase (New England Biolabs, M0495L) and 2 U of *HpaII* or *MspI* (New England Biolabs) in 20 mM Tris-HCl pH 8.3 (Merck, 648310-M), with concentrations of 80nM each for reference-F, reference-R, reference HEX, handle-F, handle-R and target FAM oligonucleotides, and 2.5nM each for the Foligo-F and Foligo-R oligonucleotides (Supplementary Table 3 and Extended Fig.1b). Reactions were sealed (Bio-Rad MSB1001), mixed using MixMate (Eppendorf), briefly centrifuged, then thermal cycled on a Bio-Rad CFX real-time PCR detection system. Conditions were 37 °C for 15 min for restriction digestion, followed by 90 °C for 5 s then 95 °C for 2 min for initial denaturation. Preamplification was performed for 10 cycles of 90 °C for 5 s, 95 °C for 15 s, 60 °C for 1 min, and 68 °C for 20 s. Quantitative PCR was then carried out for 40 cycles of 95 °C for 15 s, 60 °C for 1 min, and 68 °C for 20 s with fluorescent signal measured at the end of the 60 °C step.

#### Single-cell L1-ES-qPCR (outside of SCIMETAR-seq)

Unless otherwise stated, all reagents were added to 96-well plates using a Mantis liquid handling system (Formulatrix). After every reagent addition, plates were sealed (Thermo AB0558), vortexed briefly (1650 rpm for 30s, Eppendorf Mixmate) and centrifuged briefly (500g) before proceeding. Immediately after thawing cells single-sorted into 4µL/well lysis buffer (0.1% Tween 20, 0.1 mM EDTA, 2.5 mM Tris-HCl pH 7), 3µL/well of protease solution (0.35mg/mL Qiagen protease (Qiagen 19155) in 44 mM Tris-HCl ph 8.3 (Merck 648310-M), 1.07 mM EDTA) was added. Digestion of proteins was at 50°C for 90 min, then stopped at 75°C for 30 min. Next, 3 µL/well of restriction digestion solution (50 mM Tris-HCl pH 8.3, 250 mM KCl and 35 mM MgCl₂ containing 5U/well of either *HpaII* (NEB R0171L) or *MspI* (NEB R0106S)) was added and sealed mixed plates were incubated at 37 °C for 90 min, then 80 °C for 20 min. Finally, 10µL/well of single cell qPCR master mix (20 mM Tris-HCl pH 8.3, 100 mM KCl (Merck 60142), 400 µM dNTP, 0.5U Hot Start Taq DNA polymerase, 160 nM each for reference-F, reference-R, reference HEX, handle-F, handle-R and target FAM oligonucleotides, and 5 nM each for the Foligo-F and Foligo-R oligonucleotides (Supplementary Table 3 and Extended Fig.1b) was added. After plates were sealed and mixed, PCR conditions consisted of 95°C for 2 min, 10 preamplification cycles of 90°C for 5 s, 95°C for 15 s, 60°C for 1 min, and 68°C for 20 s, then 40 qPCR cycles of 95°C for 15 s, 60°C for 1 min and 68°C for 20 s with fluorescent signal measured at the end of the 60 °C step.

#### Integration of L1-ES-qPCR into multi-omics SCIMETAR-seq

Immediately after single-cell lysis, synthesis of full-length 1^st^ strand cDNA ^42, 43^ occurs (Fig.1a(ii)). Chromatin is deproteinated (Fig.1a(iii)) then digested with *Hpa*II (Fig.1a(iv)); cDNA is protected from *Hpa*II digestion (Extended Fig.1j, k). Full-length cDNA plus target amplicons (genomic and/or cDNA) plus L1-ES-qPCR amplicons are then multiplex pre-amplified (Fig.1a(v)). Finally, reactions are split 3-ways to complete (1) whole transcriptome cDNA libraries, (2) nested amplification, well-barcoding and plate-indexing of target amplicons, and (3) FAM/HEX-tracked L1 qPCR reactions (Fig.1a(vi)).

### SCIMETAR-seq multi-omic workflow

#### 1. Reverse Transcription, protein digestion, restriction digestion and

#### preamplification

Unless otherwise stated, all reagents were added *via* automation and wells sealed and vortexed as described above before proceeding to temperature cycling. Frozen plates containing single cells in 4µL/well lysis buffer (2.5mM Tris-HCl (Thermo AM9850G) pH 7.0, 0.1% Tween-20 (Merck P9416), 0.1mM EDTA (Merck 03690), 0.53U/µL recombinant RNase inhibitor (RRI, New England Biolabs (NEB) M0314L), 6.25% PEG 8000 (Merck P1458), 0.625 mM dNTPs (NEB N0446S) and 0.625 µM oligo-dT (Supplementary Table 3)) were centrifuged at 300g for 1 min at 4 °C, then incubated at 70 °C for 3 min to complete cell lysis and anneal mRNA to oligo-dT. RT master mix (1µL/well) was then added. RT master mix comprised 25mM Tris-HCl pH ∼8.3, 30mM NaCl (Thermo AM9760G), 1mM GTP (Thermo R1461), 2.5mM MgCl₂ (Merck M1028), 8mM dithiothreitol (Thermo 70726), 0.5U/µL recombinant RNase inhibitor (NEB M0314L), 2 µM template switching oligonucleotide (TSO, Supplementary Table 3) and 2U/µL Maxima H Minus Reverse Transcriptase (Thermo EP0752), in nuclease-free water. Reverse transcription (RT) conditions were 42 °C for 90 min, 10 cycles of 42 °C for 2 min and 50 °C for 2 min, then 85 °C for 5 min. To deplete excess oligonucleotides, 0.5µL/well of thermolabile Exonuclease I (ExoI, 20U/µL, NEB M0568L) was added and plates were incubated at 37 °C for 20 min then 80 °C for 5 min. To digest proteins, 2 µL Qiagen protease (Qiagen 19155, 2.1mg/mL in 7mM EDTA pH8.0) was added, then plates were heated at 50 °C for 90 min, followed by protease inactivation at 75 °C for 30 min. For digestion at CCGG sites, 3µL/well of RE master mix (33 mM Tris-HCl pH 8.3, 167 mM KCl, 23 mM MgCl₂ and 5 U/well of either *HpaII* (all wells in Fig 1 experiments; 92% wells in Fig 2 experiments) or *MspI* (8% wells in Fig 2 experiments)) was added. Restriction digestion was at 37 °C for 90 min, then inactivated at 80 °C for 20 min. Multiplex preamplification primers were added next in 10µL/well preamp cocktail (0.8 U/well Platinum II *Taq* DNA polymerase (Thermo 14966005), 1.5x Platinum II PCR buffer (Thermo), 0.2mM dNTPs (Thermo R0181), 50 ng ET single-stranded DNA binding protein (NEB M2401S), 10 nM each of L1 reference-F and reference-R primers, 5 nM each of L1 Foligo-F and Foligo-R, 20 nM each of L1 handle-F and handle-R primers, 40 nM each of forward and reverse “outer” primers targeting *TET2* and *SRSF2* amplicons, and 200 nM each of cDNA library forward and reverse primers). Thermal cycling was at 94 °C for 2 min, then 14 cycles of 94 °C for 20 s, 60 °C for 30 s and 68 °C for 6 min, plus a final extension at 68 °C for 5 min. To digest excess oligonucleotides, 1 µL (20U)/well thermolabile ExoI was added, then plates incubated at 37 °C for 20 min, followed by ExoI inactivation at 80 °C for 5 min. Sealed plates were stored at -20°C.

#### 2. Full-length cDNA library

To create full length cDNA libraries, 20 µL/well of cDNA master amplification mix (0.94x Platinum II PCR buffer, 2.5U/well LongAmp Hot Start Taq DNA polymerase (NEB M0534L), 125 µM dNTPs and 250 nM each of cDNA forward and reverse primers) was dispensed into fresh plates (Mantis liquid handling system), then 5 µL of pre-amp reaction (above) was added as template (Viaflo liquid handling system, Integra Biosciences). cDNA amplification involved initial denaturation at 94 °C for 2 min, then 8 cycles of 94 °C for 20 s, 60 °C for 30 s and 65 °C for 6 min, plus a final extension at 65 °C for 10 min. PCR products (10 µL aliquot) were purified using AMPure XP beads (Beckman Coulter, A63881) at a 0.6:1 bead-to-sample ratio and eluted in 10 µL nuclease-free water using Viaflo liquid handling system, then tagmented using a Nextera XT DNA Library Preparation Kit (Illumina, FC-131-1096). For each tagment reaction, 1.5 µL/well of tagmentation mix (1 µL TD reaction buffer plus 0.5 µL Tn5) was pre-dispensed into 96-well plates (MANTIS liquid handling system), then 0.5 µL cDNA was added (Viaflo) and the mix incubated at 55 °C for 5 min, then neutralized by adding 0.5 µL NT buffer (MANTIS liquid handling system) and incubating at room temperature for 5 min. cDNA was well-indexed by adding 1.0 µL of dual-index primer mix (1 µM each of i5 and i7 barcoding primers, Supplementary Table 5; in a pairing unique to each well) and 1.5 µL Nextera PCR Master Mix, followed by PCR with an initial extension at 72 °C for 3 min and denaturation at 95 °C for 30 s, then 15 cycles of 95 °C for 10 s, 55 °C for 30 s and 72 °C for 30 s, plus a final extension at 72 °C for 5 min. Tagmented libraries were pooled (1 pool per plate) and purified using AMPure XP bead at a 0.55:1 bead-to-sample ratio and eluted into ½ original volume nuclease-free water. cDNA concentration in each pool was estimated using High Sensitivity D1000 ScreenTape (Agilent, 5067-5584), normalise-pooled into a single library and sequenced (150 base paired-end reads) at the Ramaciotti Centre for Genomics on a NovaSeq X Plus platform.

#### 3. Target Amplicon Library

Target amplicons were generated by nested amplification (*TET2* reactions separate from *SRSF2*) using multiplex pre-amplification products as template. Amplicon master mixes (9 µL/well of 0.5 U Hot Start *Taq* DNA polymerase, 1× Standard *Taq* Reaction Buffer, 1 mM MgCl₂, 0.2 µM dNTPs, 0.2 µM amplicon-specific forward and reverse “inner” primers, plus 1 M betaine (Sigma-Aldrich, B0300) for *SRSF2* amplicons, were dispensed into fresh 96-well plates (Mantis liquid handling), then 1µL preamplification reaction was added (Viaflo liquid handling). Initial denaturation was at 95 °C for 3 min; *TET2* reactions were then amplified for 25 cycles of 95 °C for 20 s, 60 °C for 30 s and 68 °C for 1 min; *SRSF2* reactions were amplified for 32 cycles of 95 °C for 20 s, 58 °C for 30 s and 68 °C for 1 min; all reactions were then completed by a final extension at 68 °C for 5 min. Target amplicons were then barcoded using reaction conditions identical to above (i.e. 9 µL master mix plus 1 µL nested amplicon as template) using forward barcode primers to identify plate column and reverse barcode primers to identify plate row (Supplementary Table 4). *TET2* cycling conditions were 6 cycles of 95 °C for 20 s, 60 °C for 30 s and 68 °C for 1 min, followed by a final extension at 68 °C for 5 min. *SRSF2* cycling conditions were 9 cycles of 95 °C for 20 s, 58 °C for 30 s and 68 °C for 1 min, followed by a final extension at 68 °C for 5 min. Amplicons were pooled by plate then purified using AMPure XP beads at a 0.6:1 bead-to-sample ratio and quantified using High Sensitivity D1000 ScreenTape. Illumina P5 and P7 adapters and indexes were added to each pool by the Ramaciotti Centre for Genomics sequencing facility. Libraries were normalized pooled and sequenced on a MiSeq i100 Plus platform with 20% PhiX spike-in using 300 base paired-end reads.

#### 4. L1-ES-qPCR

SCIMETAR L1 ES-qPCR master mix (16 µL) was dispensed (Mantis liquid handling) into fresh 96-well plates. Master mix consisted of 18.75 mM Tris-HCl pH 8.3, 50 mM KCl, 2.5 mM MgCl₂, 0.25 µM dNTPs, 0.5 U/well Hot Start Taq DNA polymerase, 0.05 µM each of the five L1 target oligonucleotides, and 0.1 µM each of the three L1 reference oligonucleotides. Template (4 µL) was then added from multiplex primary PCR plates by Viaflo liquid handling. Thermal cycling conditions consisted of 37 °C for 2 min, 90 °C for 5 s, and 95 °C for 2 min, followed by 40 cycles of 95 °C for 15 s, 65 °C for 40 s, and 68 °C for 20 s, with a final hold at 12 °C and fluorescent signal measured at the end of the 65 °C step.

### Data Analysis

#### Statistics used for Figures

All statistics were produced in R or GraphPad Prism v10.6.1. Statistical tests used for specific figure panels are as indicated below.

*Figure 1:* (c) Adjusted *p* values from Šídák’s multiple comparisons tests. (e) Tukey’s box plots used; adjusted *p* values from Šídák’s multiple comparisons tests. (f) FDR *q* values (Benjamini, Krieger & Yekutieli) determined from multiple Fisher’s exact 2×2 (top: in cluster, out of cluster x vehicle, AZA; bottom: vehicle, AZA x cord blood, PD7151) contingency test *p* values. (g) Unadjusted *p* values from Fisher’s 2×2 exact tests (left: CMP, not CMP x *SRSF2*-wt, *SRSF2*-mut; right: cluster-2, not cluster-2 x *SRSF2*-wt, *SRSF2*-mut).

*Figure 2:* (b): *p*-values for Welch’s t-tests. (d) Nonparametric Spearman correlation r and p values, plus *p* values for non-zero slope from simple linear regression. (f-g) FDR *q* values (Benjamini, Krieger & Yekutieli) determined from multiple Fisher’s exact 2×2 (in cluster, out of cluster x D1, D8) contingency test *p* values. (h) Adjusted *p* values for nonparametric Dunn’s multiple comparisons tests, always comparing to C4D1 cluster-1 cells. (i-j) Tukey box plots overlaid. Adjusted *p* values are from comparing every group with every other group (Dunnett’s multiple comparisons test). (k) FDR *q* values (Benjamini, Krieger & Yekutieli) from multiple Fisher’s exact 2×2 (in mt cluster, out of mt cluster x D1, D8) contingency test *p* values. (l) FDR *q* values (Benjamini, Krieger & Yekutieli) from multiple Fisher’s exact 2×5 (in mt cluster, out of mt cluster x RNAseq cluster-0, -1, -2, -3, -4) contingency test *p* values, performed separately for D1 and for D8.

*Extended Fig.1:* (h) Adjusted *p* values from Tukey’s multiple comparisons tests. (i) Adjusted *p* values from Holm-Šídák’s multiple comparisons tests. No test was applied to compare *Hpa*II with *Msp*I data.

*Extended Fig.2:* (e) Adjusted *p* values for nonparametric Dunn’s multiple comparisons of every column with every other column.

*Extended Fig.3:* (d) FDR *q* values (Benjamini, Krieger & Yekutieli) determined from multiple within immunophenotype Fisher’s exact 2×2 (vehicle, AZA x cord blood, PD7151) contingency test *p* values. (f) Adjusted *p* values for within immunophenotype vehicle versus AZA Šídák’s multiple comparisons tests. (k) Unadjusted *p* value from Fisher’s 2×2 exact test (CMP, not CMP x *SRSF2*-wt, *SRSF2*-mut).

*Extended Fig. 4:* (a) FDR *q* values (Benjamini, Krieger & Yekutieli) determined from multiple within immunophenotype Fisher’s exact 2×2 (immunophenotype, not immunophenotype x C4D1, C4D8) contingency test *p* values. (c) Nonparametric Spearman correlation r and p values, plus *p* values for non-zero slope from simple linear regression. (d) Welch’s t test *p* for all HSC immunophenotype versus all other cells.

*Extended Fig. 5:* (a) Adjusted *p* values for nonparametric Dunn’s multiple comparisons of every column with every other column. (b) Mann-Whitney test *p* value.

#### L1-ES-qPCR Data Analysis

qPCR data were analyzed using CFX Maestro software (v2.3; Bio-Rad) to determine HEX and FAM Ct values. Relative demethylation was calculated using the equations in Extended Fig.1f.

#### Cell type assignment on index-sorted cells

Fluorescence intensity data recorded during index sorting were analysed as previously described 24.

#### Single-cell RNA-seq Alignment and Quantification

Code used to perform alignment and analysis can be found at https://github.com/alexyfyf/SCIMETAR. In brief, raw FASTQ files from single-cell RNA-seq were aligned to the GRCh38 reference genome using STAR (v2.7.11b) in 2-pass mode (--twopassMode Basic) ^44^. A splice junction overhang of 150 bp (--sjdbOverhang 150) was used and we also set --outSAMmultNmax 1 and --outFilterMultimapNmax 50 as used by zUMIs pipeline ^45^. Gene expression quantification was performed on the aligned BAM files using featureCounts (v2.0.8) with Gencode v44 annotation, parameters -p (for paired-end data) and -s 0 (unstranded) ^46^. The output was a gene-by-cell count matrix. Code for post-alignment analysis can be found at https://github.com/alexyfyf/SCIMETAR_seq_paper_analysis.

#### Targeted Amplicon Sequencing Processing

##### Demultiplexing

Raw amplicon data was demultiplexed using Matchbox (v0.1.0) ^47^. Reads were assigned to samples based on unique combinations of forward and reverse barcodes identified at the 5’ ends of Read 1 and the 3’ ends of Read 2 (reverse complement), allowing for a 1 bp mismatch.

##### Genotyping

To genotype each sequencing fragments, we perform mutation calling separately in Read1 (R1) and Read2 reverse complement (R2rc) in 2 rounds. In the first round, we look for an exact 5bp match centred around the mutation site (Supplementary Table 6) in both R1 and the R2rc for TET2 [exon3], TET2 [exon6] and SRSF2. Any reads that didn’t match in round 1 are searched again specifically for SRSF2, using a 10bp flanking sequences and allowing for a 1bp mismatch.

For each read, we then genotype them as (1) TET2 [exon3] WT or MUT only if the sequence was found in R1 alone, or in both R1 and R2rc. If it was only found in R2rc, it is discarded. (2) TET2 [exon6] WT or MUT only if the sequence was found in R2rc alone, or in both R1 and R2rc. (3) SRSF2 WT or MUT if the sequence was found in R1 alone, both R1 and R2rc, or for in vivo data only if it was recovered during the less stringent Round 2 search.

To determine the genotype of each cell for a particular variant, cells with less than 20 or 10 genotyped reads, for in vitro and in vivo respectively, are labelled “unidentifiable”. We then calculated the ratio of mutant reads / (mutant + wildtype reads), and if the ratio is > 0.99, the cell is labelled “mut/mut”, if the ratio is < 0.05, the cell is labelled “wt/wt”, and rest labelled “mut/wt”.

#### Single-cell RNA-seq Data Analysis

The analysis was performed using R version 4.4.2. The gene expression count matrix was loaded into the Seurat R package (v5.1.0) and RPKM normalized ^48^. Cells were filtered to retain high-quality cells. Empty wells, or cells called as dead based on FACS index data, were excluded. Remaining cells were included if they had more than 1000, 2000 or 1000 detected features (for MDSL cell line, in vitro and C4 in vivo datasets respectively). Raw counts were normalized and variance stabilized using the vst selection method in Seurat to identify the top 2,000 variable features. Data was scaled, and Principal Component Analysis (PCA) was performed on the variable features. The top 20 (10 for MDSL) principal components were used for downstream clustering and dimensionality reduction. A Shared Nearest Neighbor (SNN) graph was constructed with k=50 (30 for MDSL) neighbors, and clusters were identified using the Louvain algorithm with a resolution of 1.0 (0.8 for MDSL). Uniform Manifold Approximation and Projection (UMAP) was calculated on the same PCA dimensions to visualize the data in two dimensions. Cell cycle scoring was performed to assess the G2M and S phase scores for each cell.

#### Cell Type Annotation

Automated annotation was performed using a reference dataset from a reference paper ^26^ using Azimuth (v0.5.0) to align with established haematopoietic hierarchies ^26, 49^. Cluster-specific marker genes were identified using the FindAllMarkers function in Seurat with a Wilcoxon Rank Sum test. The resulting markers were compared against canonical markers for validation (e.g., AVP, CRHBP, DLK1 for HSC/MPP; MPO, CDK6 for Neu1; EBM markers). Pathway analysis of marker genes was performed using clusterProfiler package and Gene Ontology database ^50–52^.

#### Genotype-Phenotype Integration

Targeted amplicon sequencing data for *SRSF2, TET2* exon 3*, and TET2* exon 6 mutations was integrated with the single-cell transcriptomic data. Genotyping results were matched to single cells using plate and well information. Cells were categorized based on their mutational status (e.g., WT/WT, Mut/WT, Mut/Mut). We correlated genotypes with cell phenotypes by constructing Sankey/Alluvial plots to visualize the flow between FACS immunophenotypes, Seurat clusters, and reference-based annotations across different experimental conditions (e.g., DMSO vs. AZA treatment).

#### LINE-1 qPCR Analysis

Quality control was performed by analyzing the distribution of reference gene Ct values (HEX_Ref) across plates and experimental groups. Outlier wells were identified and removed using a Median Absolute Deviation (MAD) approach, retaining wells where HEX_Ref was within the median ± 2 * MAD (group-wise for in vitro and plate-wise for in vivo). For the past wells, dCT was calculated as Ct (FAM_Tar) – Ct (Hex_Ref). For the in vivo C4 dataset, we use *MspI* wells as internal control to normalize across plates and remove batch effects, limma::removeBatchEffect was used, and we used plate as batch and *MspI* wells as design matrix^53^.

#### Mitochondrial variation

Mitochondrial allele counts were first extracted from scRNA-seq using cellSNP-lite^54^. For mitochondrial (mt) SNV clusters, informative mt variants (with confident non-zero allele frequency) were first selected using MQuad^55^ with default parameters, and then cells with similar mt genotypes were clustered into lineages using SNPmanifold^56^ with default parameters and k-means clustering.

### Instrumentation

Details of the instrumentation used to support the single cell workflow is provided in Supplementary Table 7.

---

**Extended Figure 1:**
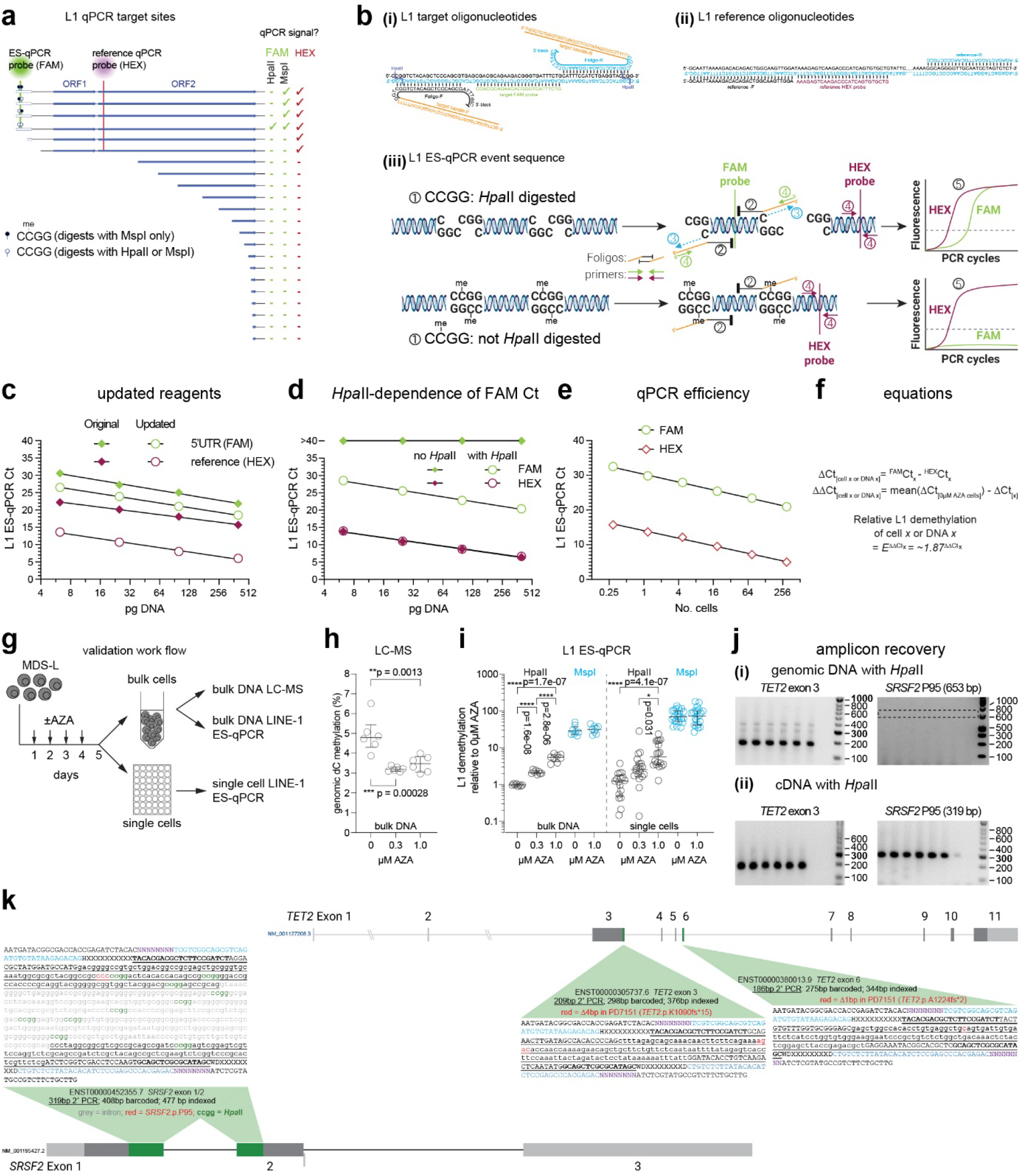
Development and validation of a single-cell LINE1 ES-qPCR assay to assess genome-wide CpG methylation. **(a)** Schematic showing LINE1 (L1) regions targeted by qPCR; most L1 elements in the human genome are 5’-truncated ^20^. **(b)** Outline of L1-ES-qPCR showing, **(i)** L1 5’UTR ES-qPCR oligonucleotides, **(ii)** L1 5’ORF2 reference oligonucleotides, **(iii)** reaction sequence of L1-ES-qPCR (^①^ Restriction digestion; ^②^ Annealing of L1-specific foligos; ^③^ extension of genomic 3’-ends created by restriction digestion – *if* the 3’-ends are annealed to a foligo; ^④^ copying of (green) foligo-tagged templates or (red) reference templates by PCR; ^⑤^ detection of products by quenched probes. **(c)** Sensitivity of original (solid symbols; Rand *et al.* (2010) *BioTechniques*) ^17^ versus our updated (open symbols) L1 ES-qPCR reagents. **(d)** *HpaII-*digestion-dependence of L1 ES-qPCR FAM signals. **(e)** Ct plotted as function of log_2_[DNA]); used for PCR efficiency calculations. **(f)** ΔCt and ΔΔCt definitions (top), and relative L1 demethylation estimations (bottom). **(g)** Workflow to compare L1 ES-qPCR to LC-MS. Cultured MDS-L cells were treated daily with 0, 0.3 or 1.0 µM AZA for 4 consecutive days, then harvested on the fifth day for bulk cell DNA extraction (top) or sorted as single viable cells into lysis solution (bottom). **(h)** Genome-wide percent cytosine methylation in the AZA-treated MDS-L DNA, measured by LC-MS. n=6 technical replicates, data sourced from ^27^; *p:* Tukey’s multiple comparisons tests. **(i)** Relative L1 demethylation (normalized to 0µM AZA treatment) as determined by ES-qPCR from (left) bulk DNA or (right) single MDS-L cells; *p:* Holm-Šídák’s multiple comparisons tests. No test was applied to compare *Hpa*II with *Msp*I data. **(j)** Examples of single cell targeted amplicon recoveries from **(i)** *Hpa*II-digested genomic gDNA or **(ii)** *Hpa*II-digested first strand cDNA. **(k)** Detail of complete target amplicons for *TET2* (upper) and *SRSF2* (lower): (green) *Hpa*II sites, (red) target mutations sites, (purple N) I5 or i7 index, (X…X) well barcodes, (<u>underlined</u>) exonic sequence.

**Extended Figure 2:**
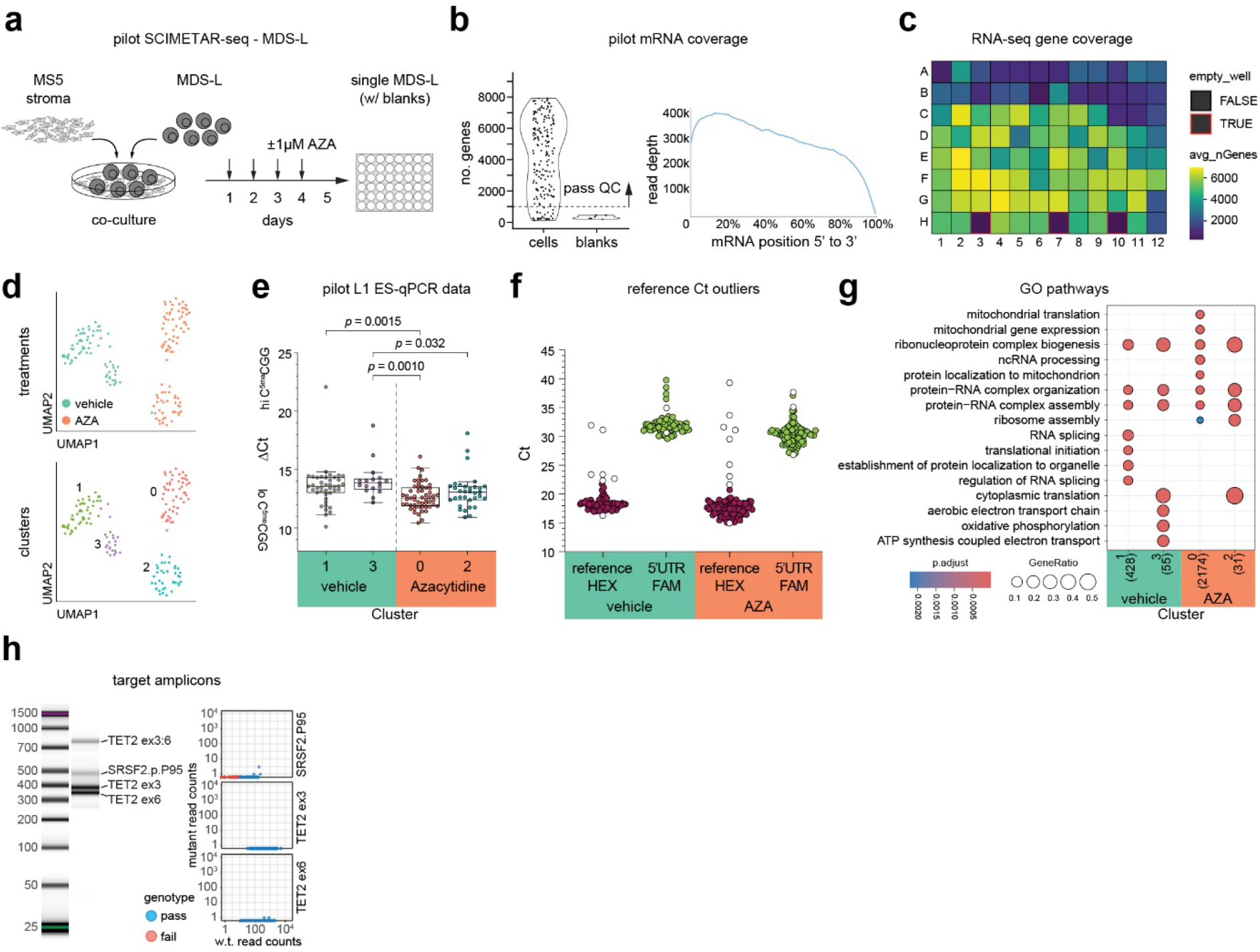
Pilot implementation of SCIMETAR-seq in the MDS-L cell line. **(a)** Schematic showing in vitro AZA treatment of MDS-L co-cultured with murine MS5 stromal cells. **(b)** Genes recovered per cell (left) and normalized 5’- to 3’ transcript coverage (right). **(c)** Heatmap showing average gene coverage per well (1 vehicle plate; 1 AZA plate). **(d)** UMAP embedding of MDS-L cells colored by treatment or cluster. **(e)** *Δ*Ct (Ct_FAM_ - Ct_HEX_) distribution for RNA-seq clusters; Tukey box plots; *p:* Dunn’s multiple (all way) comparisons. **(f)** Rationale for filtering L1 ES-qPCR outliers (see Methods): all single cells contain the same number of L1 reference DNA templates but vary in L1 methylation levels. Reference (HEX) Ct falling outside 2x mean absolute deviation were excluded as outliers that failed reference qPCR (plotted as white circles). Most, but not all HEX Ct outliers failed to produce a FAM Ct value or were FAM Ct outliers. **(g)** Gene ontology (GO) pathways significantly enriched in each MDS-L cluster. **(h)** Bioanalyzer trace of SCIMETAR target amplicon library (left) and plots of w.t. versus mutant reads for each target (right).

**Extended Figure 3:**
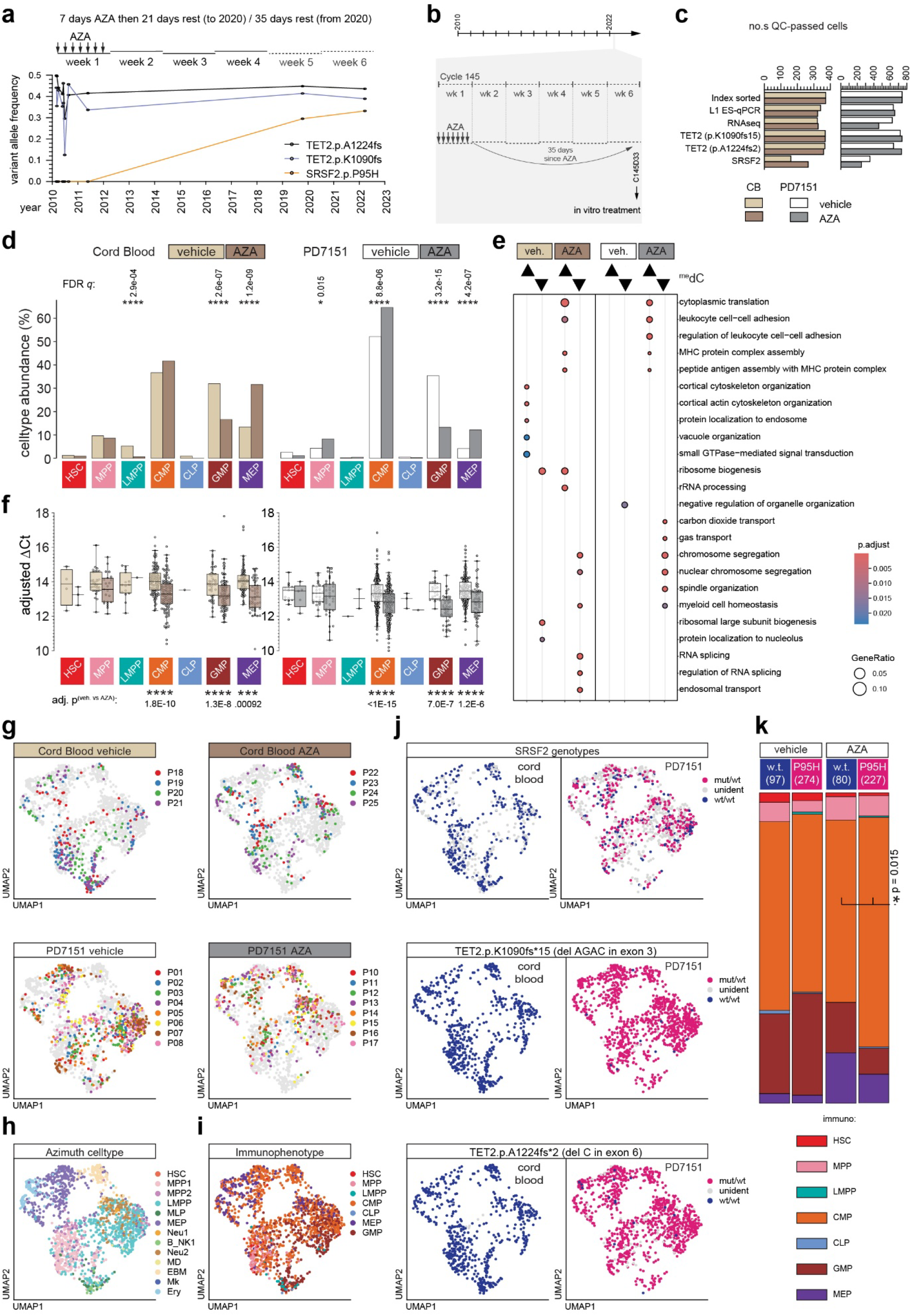
SCIMETAR-seq analysis of *in vitro* AZA-treated cells. **(a)** Treatment history and longitudinal variant allele frequencies for long-term AZA responding patient PD7151. **(b)** Schematic showing treatment context of sample used for *in vitro* AZA treatment. Bone marrow aspirate was collected immediately prior to the start of clinical AZA treatment cycle 146, 33 days after the most recent administration of AZA. **(c)** Total number of cells that passed QC for each SCIMETAR-seq component. Twice as many PD7151 as CB cells were index sorted. **(d)** Relative abundance per immunophenotype; *q:* FDR for multiple 2×2 contingency tests of vehicle vs AZA. **(e)** Pathway enrichment for genes differentially expressed between most (▴) and least (▾) methylated cells. **(f)** Per-immunophenotype single-cell adjusted ΔCt; *p:* Šídák’s multiple vehicle vs AZA comparisons. (**g-j**) UMAP embeddings coloured by **(g)** sort plate, **(h)** transcriptomic identity **(i)** immunophenotype, **(j)** *SRSF2*, *TET* exon 3 or *TET2* exon 6 genotypes for (left) CB and (right) PD7151 cells. **(k)** Relative abundance of PD7151 immunophenotypes per treatment and *SRSF2* genotype; *p:* 2×2 (CMP, not CMP x *SRSF2-*wt, -mut) contingency test for the impact of genotype on CMP frequency.

**Extended Figure 4:**
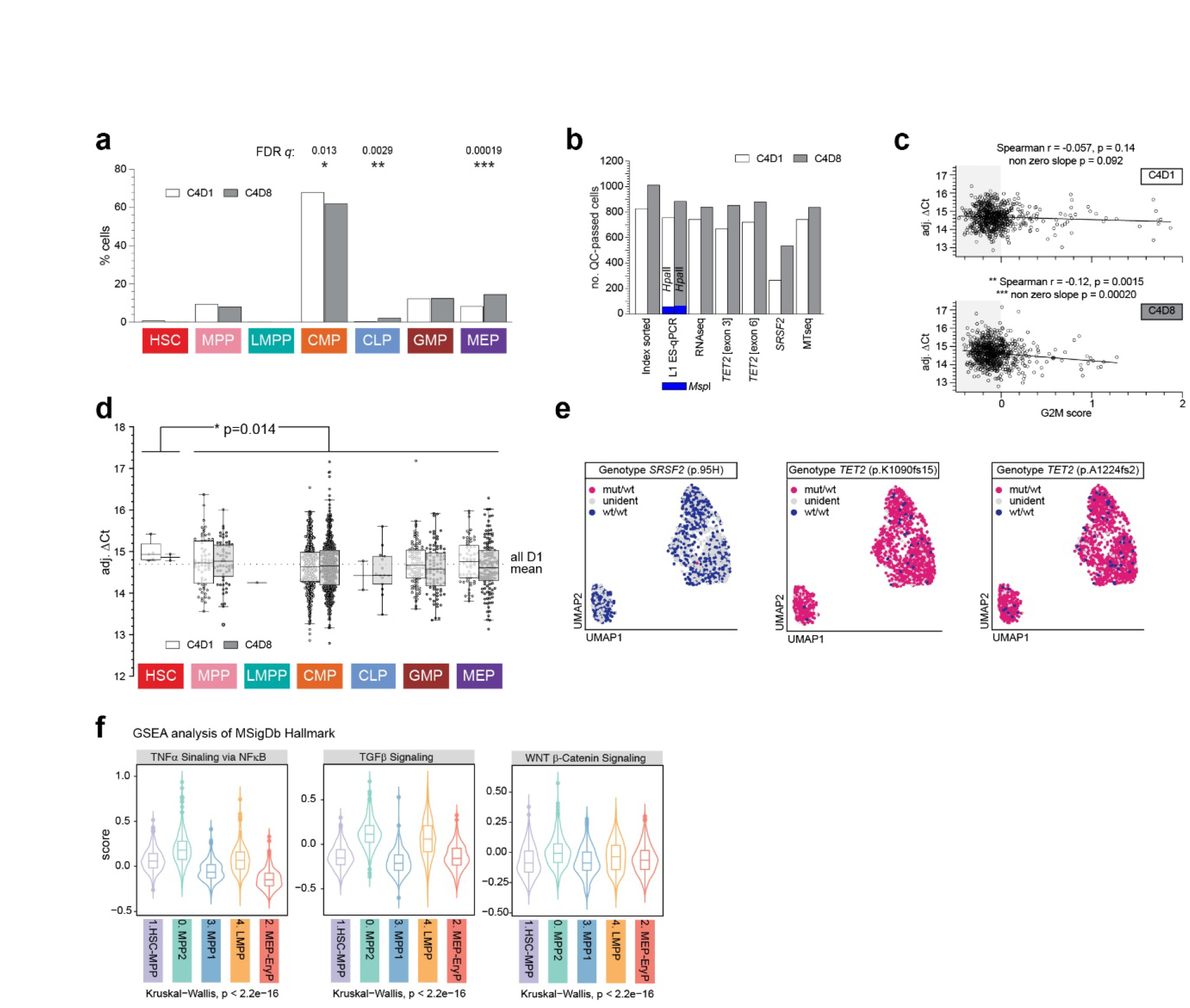
SCIMETAR-seq analysis of *in vivo* AZA-treated cells. **(a)** Relative abundance per immunophenotype; *q:* FDR for multiple 2×2 (immunophenotype, not immunophenotype x D1, D8) contingency tests. **(b)** Total number of cells that passed QC for each SCIMETAR-seq component, including sufficient mitochondrial reads for SNPmanifold (mt genotypes). **(c)** ΔCt compared to transcriptomic G2M-score, grey shading indicates cells not in S-phase; Spearman correlation and linear regression. **(d)** Per-immunophenotype single-cell adjusted ΔCt; *p:* Welch’s t-test comparing all HSC immunophenotype to all other cells, regardless of timepoint. **(e)** UMAP embeddings coloured by *SRSF2*, *TET* p.K1090fs*15 (exon 3) or *TET2* p.A1224fs*2 (exon 6) genotypes. **(f)** Hallmark signaling pathways significantly enriched in cluster-0 cells.

**Extended Figure 5:**
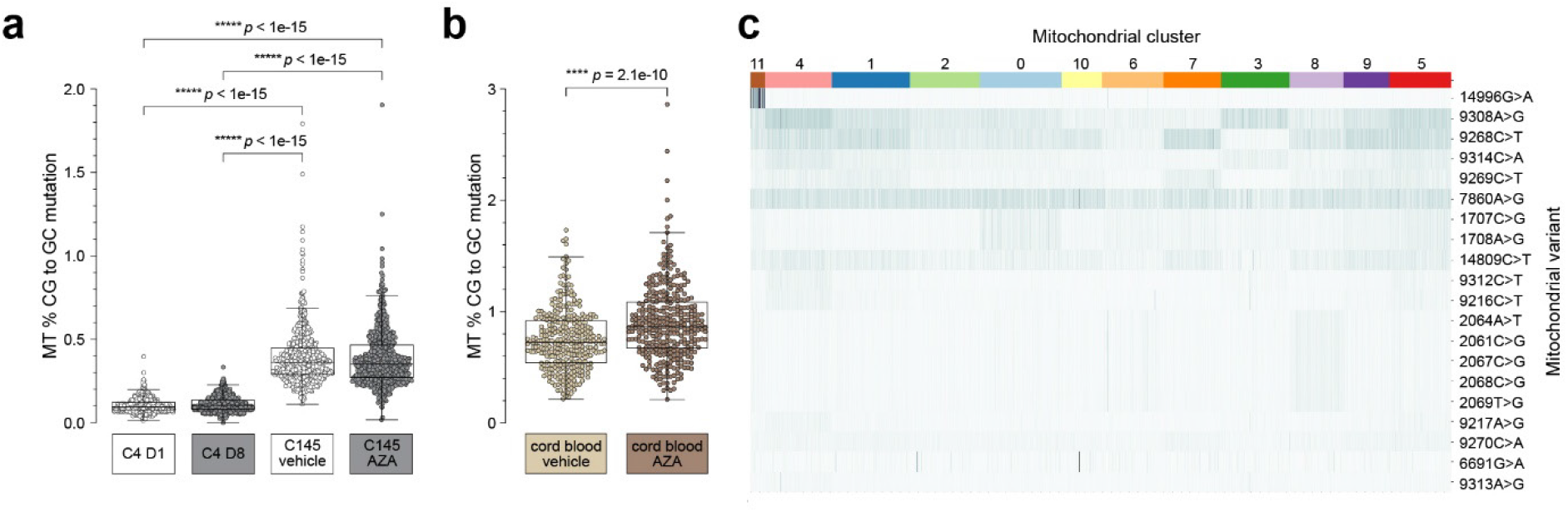
**(a)** Frequencies of C•G to G•C mutation in PD7151 HSPC mt grouped by clinical cycle number (C4 or C145) and *in vivo* treatment day (D1 or D8) or *in vitro* treatment type (vehicle or AZA); *p:* multiple nonparametric all-ways comparisons. **(b)** Frequencies of C•G to G•C mutation in cord blood HSPC mt grouped by *in vitro* treatment type; *p:* Mann-Whitney test. **(c)** SNPmanifold clustering into lineages, of the most abundant mitochondrial variants in PD7151 C4 cells

## Notes

### Competing Interest Statement

The authors have declared no competing interest.

### Summary of Updates

Manuscript and figures have been substantially reworked to improve clarity and improve statistical analyses. Additional data relating to AZA-induced mutations in mitochondrial DNA is now included.

